# Dedifferentiating germ cells regain stem-cell specific polarity checkpoint prior to niche reentry

**DOI:** 10.1101/2023.04.26.538507

**Authors:** Muhammed Burak Bener, Autumn Twillie, Rakshan Chadha, Naiya Patel, Mayu Inaba

## Abstract

In the Drosophila germline stem cell system, maintenance of the stem cell pool requires “dedifferentiation”, in which differentiating cells reattach to the niche and reacquire stem cell properties. However, the mechanism of dedifferentiation remains poorly understood. Here, using long-term live imaging, we show that dedifferentiated cells immediately re-enter mitosis with correct spindle orientation after reattachment to the niche. Analysis of cell cycle markers revealed that these dedifferentiating cells are all in G2 phase. In addition, we found that the observed G2 block during dedifferentiation likely corresponds to a centrosome orientation checkpoint (COC), a previously reported polarity checkpoint. We show that re-activation of a COC is likely required for the dedifferentiation thus ensuring asymmetric division even in dedifferentiated stem cells. Taken together, our study demonstrates the remarkable ability of dedifferentiating cells to reacquire the ability to divide asymmetrically.

## Introduction

Tissue stem cells play a vital role for tissue maintenance by continuously producing differentiating cells. Maintenance of the number of stem cells throughout a lifespan is essential for multicellular organisms [1]. The loss of stem cells underlies tissue degeneration in both physiological and pathological conditions.

Stem cells are mainly maintained by asymmetric division in which each cell division results in self-renewal and differentiation. However, stem cells can be lost from the tissue due to the lifetime of individual cells [2, 3] or also under conditions such as injury, and metabolic stress [4, 5]. Lost stem cells can be replenished through two mechanisms, a symmetric division, or a dedifferentiation [6-10]. Dedifferentiation involves a differentiating or differentiated cell reverting to a stem cell-like state and is required for maintenance of GSC pool in the testis (Figure 1A) [8, 10, 11]. Dedifferentiation has been observed in various stem cell systems across both vertebrates and invertebrates, postulating that it is a conserved mechanism for stem cell maintenance [3, 6, 10, 12-14]. However, due to an anatomical complexity, the mechanism of dedifferentiation has been poorly understood in other stem cell systems.

**Figure 1.**
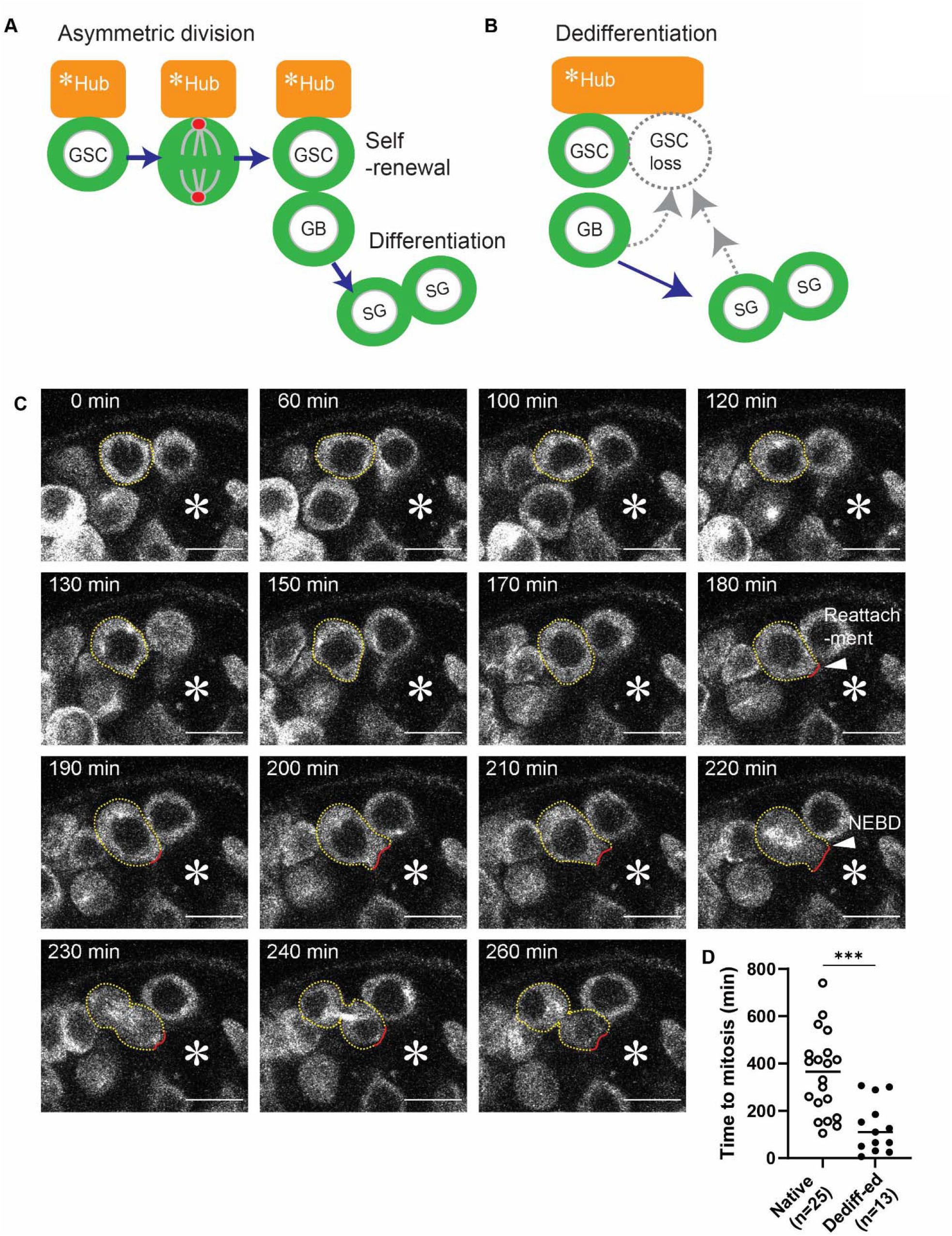
Dedifferentiated GSCs divide immediately upon niche reattachment. **A**) Schematic of asymmetric division of a GSC. GSC forms oriented mitotic spindle perpendicular to the hub-GSC interface. Spindle orientation during mitosis results in retaining one daughter near the hub as a GSC (self-renewal) and displacing the other daughter cell away from the hub as a gonialblast (GB, differentiation). Then, the GB divides to become spermatogonia (SGs). *Hub=niche. **B**) Dedifferentiation (see text) is essential for maintenance of GSCs in Drosophila testicular niche. **C**) Representative time-lapse live images of a dedifferentiating gonialblast (GB). Germ cells are visualized by expression of UAS-αTubGFP under the control of nosGal4 driver. A dedifferentiating germ cell is encircled by yellow dotted lines. The interface of dedifferentiated GSC and hub cell is marked by red lines. NEBD=nuclear envelope breakdown. **D**) The time taken between hub-reattachment and 1^st^ mitotic division (NEBD) for all recorded dedifferention events listed in Table1 is plotted to the graph. Various genotypes were used (see Table1). The cases of “not divided” were not included in the graph. P-values were calculated by a 2-tailed student-t-test and provided as *** P < 0.0001. “n” indicates the number of scored GSCs. Horizontal lines in the graph represent mean values. All scale bars represent 10 μm. The hub is marked by asterisks.

The *Drosophila* testis is an ideal model system to study stem cell maintenance in vivo. In the testes, there are 8-12 germline stem cells (GSCs) located around a cluster of cells called the hub [15]. The hub serves as a niche for maintaining the stem cell population by secreting ligands [16-20]. The GSCs in the *Drosophila* testis usually divide asymmetrically, where one daughter cell remains as a stem cell, while the other daughter cell is displaced away from the hub and begins to differentiate (Figure 1A) [21]. The differentiated cell, called the gonialblast (GB), goes through four rounds of incomplete cell division to form 16-cell spermatogonia (SG) cysts. These cysts then become spermatocytes and enter meiosis [22].

Acute stem cell loss and the recovery process can be experimentally induced in *Drosophila* gonads [6, 10, 14]. When male and female GSCs are depleted from their niche, differentiating cells migrate back to the niche, and regenerate the stem cell pool [6, 10, 14]. It was also shown that under prolonged stress conditions such as starvation, stem cells leave the niche and, and during refeeding, the lost stem cells can be recovered [23, 24]. Furthermore, dedifferentiation can also be observed in normal condition, without depletion of stem cells in *Drosophila* testis [25].

In this study, we examined the process of dedifferentiation using live imaging after depletion of GSCs. We show that, when reattached to the hub, dedifferentiated cells divide promptly (<1∼2 hours) with correct spindle orientation. Furthermore, our results suggest that a centrosome orientation checkpoint (COC), a previously reported polarity checkpoint is activated even before the cells reattach to the niche. We further show that re-activation of a COC is likely required for the dedifferentiation thus ensuring asymmetric division even in dedifferentiated stem cells. Taken together, these findings demonstrate the tight regulation of asymmetric division and the remarkable capacity of dedifferentiated cells to reacquire asymmetric division in the niche.

## Results

### Niche is replenished exclusively by dedifferentiation but not by symmetric division

To examine the behavior of cells throughout the dedifferentiation, we performed long-term live imaging (∼16 hours) after depletion of GSCs from the niche. To deplete GSCs, we forcedly expressed a differentiation factor, Bam (Bag of marbles) [26] under the control of the heat-shock promoter (*hs-bam*) as previously demonstrated [6]. Then, upon withdrawal of ectopic Bam expression, differentiating germline cells return to the niche, rapidly restoring the stem cell pool [6].

After inducing ectopic Bam expression in GSCs, the average GSC number became 1.875 ± 1.59/testis (n = 16). We then let them recover for different durations and then dissected testes to start following live-imaging for 16 hours. We observed dedifferentiation events, 0.30± 0,48 cells/testis in the period of 20-36 hour after heat shock (n = 10) and 0.84± 0.37 cells/testis in the period of 32-48 hour after heat shock (n = 19). Therefore, we used the period of 32-48-hour for following experiments.

We first noticed that, even after severe loss of GSCs, no symmetric division was observed in remaining GSCs or in already dedifferentiated GSCs (Table1). 96%± 20 (n=26) of spindles of observed in the divisions of native GSCs (defined as the cells already attached to the niche when imaging was started) were oriented, which did not show significant difference from control GSCs (94%± 23, n=36, Table 2). This result indicates that replenishment of GSC do not occur through spindle misorientation or symmetric division but occurs by dedifferentiation.

To exclude the potential influence of genotypes with different fluorescent markers on the cell behavior, we used fly testes from several genetic backgrounds, none of examined genotypes predominantly show spindle misorientation after forced depletion of GSCs (Table 1).

### Dedifferentiated GSCs divide immediately upon niche reattachment

In addition to the remarkable rate of spindle orientation in native GSCs during niche replenishment, we also noticed that dedifferentiated GSCs also divide with correctly oriented spindles (96.9%± 17.4, n=33, Table 1-3, Figure 1B) with similar frequency to original GSCs (94%± 23, n=36, Table 2), (p=0.1025). Moreover, we noticed that the dedifferentiated GSCs enter mitosis promptly upon reattachment to the niche as the average time between reattachment and 1^st^ mitotic entry of dedifferentiated GSCs was 2.18 hours ± 0.43, n=13 (Figure 1B, Table 1). It is known that average cell cycle time of male GSC is approximately 12 hours [27]. Therefore, if we assume that dedifferentiation randomly occurs in any cell cycle phase and the dedifferentiating/dedifferentiated cells undergo similar cell cycle as original GSCs, the time to 1^st^ mitosis would be randomly distributed within next 12 hours, indicating that the “time taken to mitotis” in dedifferentiated GSCs is strongly biased. Indeed, native GSCs that were already present in the niche at the start of imaging underwent division randomly in next 12 hours (6.02 hours ± 0.54, n=26 in average) from starting point of imaging (Figure 1B).

The observation that dedifferentiated GSCs promptly divide upon returning to their niche suggested two possibilities; 1. Dedifferentiating cells return to the niche in biased cell cycle phase, likely in the late-G2 phase of the cell cycle, 2. Dedifferentiated GSCs has significantly shorter cell cycle time compared to original GSCs. To distinguish these possibilities, we assessed mitotic rate of dedifferentiated GSCs after experimentally induced dedifferentiation. When we observed 2^nd^ mitotic entry of dedifferentiated GSCs, the duration between 1^st^ and 2^nd^ mitoses was consistent with normal GSC’ s (Dedifferentiated GSCs:9.9 hours ± 0.4, n=2, Native GSCs:12.9 hours ± 0.85, n=4). This indicated that dedifferentiating cells have similar mitotic rate to original GSCs. Therefore, we concluded that the dedifferentiating cells are likely posed in late G2-phase, and begin mitosis at the observed timing.

### Dedifferentiating germ cells are in G2 phase

To directly confirm the cell cycle phase of dedifferentiating cells, we next conducted live imaging using testes expressing Fly-FUCCI cell cycle marker. We found that all GSCs are actively dividing (Table 3) with their cell cycle duration approximately 12 hours (12.9 hours ± 0.85, n=4). After GSC depletion, we observed 15 dedifferentiating cells and all of them were marked in G2 phase and immediately divide after reattachment, further confirming the biased cell cycle phase in dedifferentiating cells (Table 3, Figure 2A, B).

**Figure 2.**
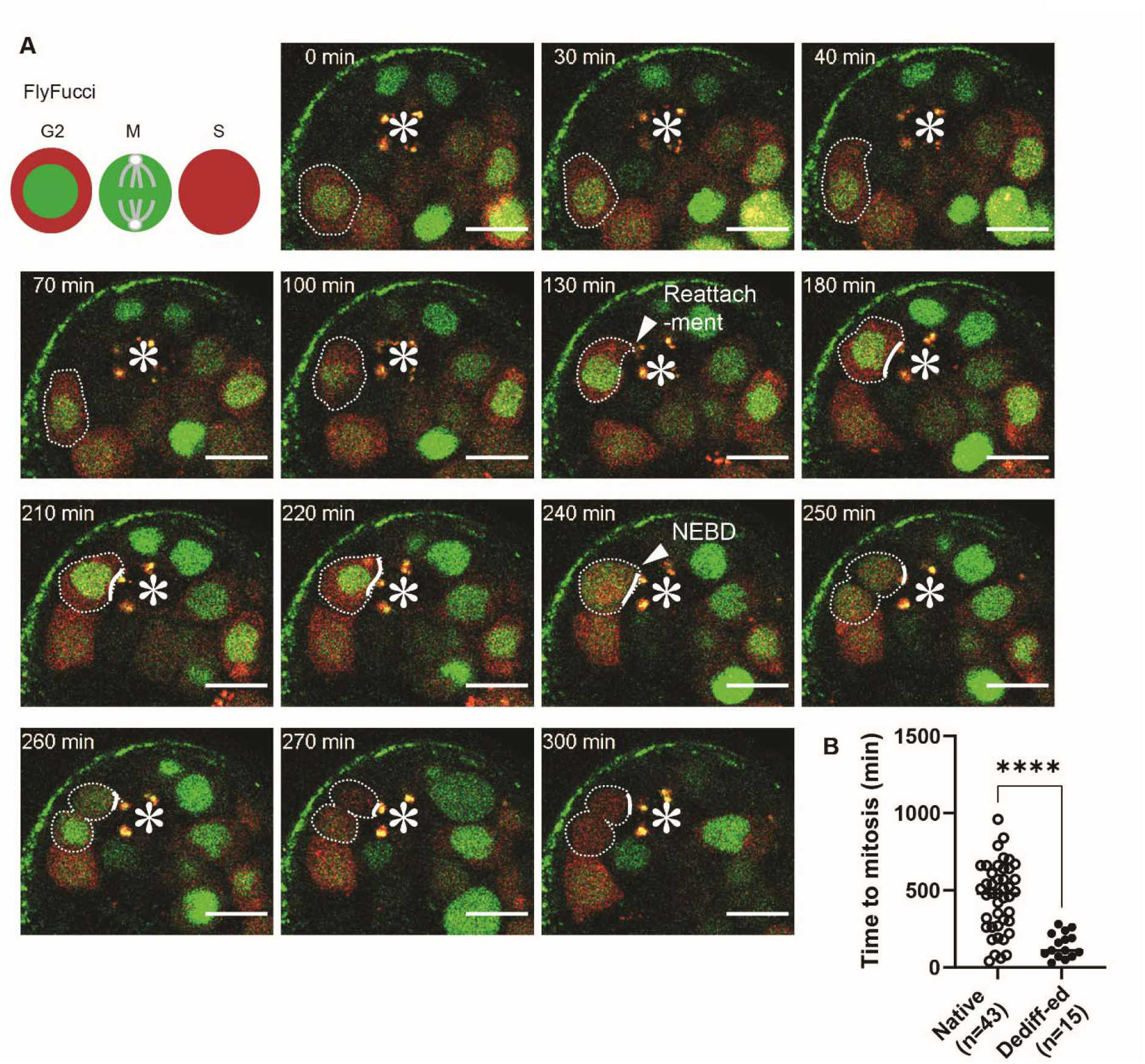
Dedifferentiating germ cells are in G2 phase. **A**) Representative time-lapse live images of a dedifferentiating gonialblast (GB) in fly-FUCCI expressing testes under the control of nosGal4 driver. GFP-E2F1-230 (green) is positive in G2-phase and negative in S-phase cells. mRFP1.CycB.1-266 (red) is positive in G2 and S phase and negative in M-phase and G1 phase. Note that G1 phase was not detectable. A dedifferentiating germ cell is encircled by white dotted lines. The interface of dedifferentiated GSC and hub cell is marked by white solid lines. NEBD=nuclear envelope break down. **B**) The time taken between hub-reattachment and 1^st^ mitotic division (NEBD) for all recorded dedifferention events listed in Table S1 is plotted to the graph. The cases of “not divided” were not included in the graph. P-values were calculated by a 2-tailed student-t-test and provided as *** P < 0.0001. “n” indicates the number of scored GSCs. Horizontal lines in the graph represent mean values. All scale bars represent 10 μm. The hub is marked by asterisks.

### Centrosome orientation checkpoint is activated in dedifferentiating germ cells

During the S-phase, GSC-GB pair has prolonged cytoplasmic connection via fusomes and the GBs are in G2 phase after they are pinched off from the GSC mother [28]. Therefore, there is the possibility that the observed G2 bias of the cell cycle is caused by the cell-type bias. To test this possibility, we sought to examine if dedifferentiating cells are “arrested” in specific cell cycle phase or if they are simply in regular G2 phase.

Potential G2 arrest of the cell cycle is reminiscent to a previously identified polarity checkpoint, centrosome orientation checkpoint (COC) (Figure 3A) [9, 29, 30]. In the male GSCs, ACD is achieved by the stereotypical positioning of centrosomes during interphase [21, 31]. The COC monitors the centrosome orientation and arrests the cell cycle at late G2 phase if centrosomes are misoriented in the GSCs, hence, ensuring the ACD [9, 29].

**Figure 3.**
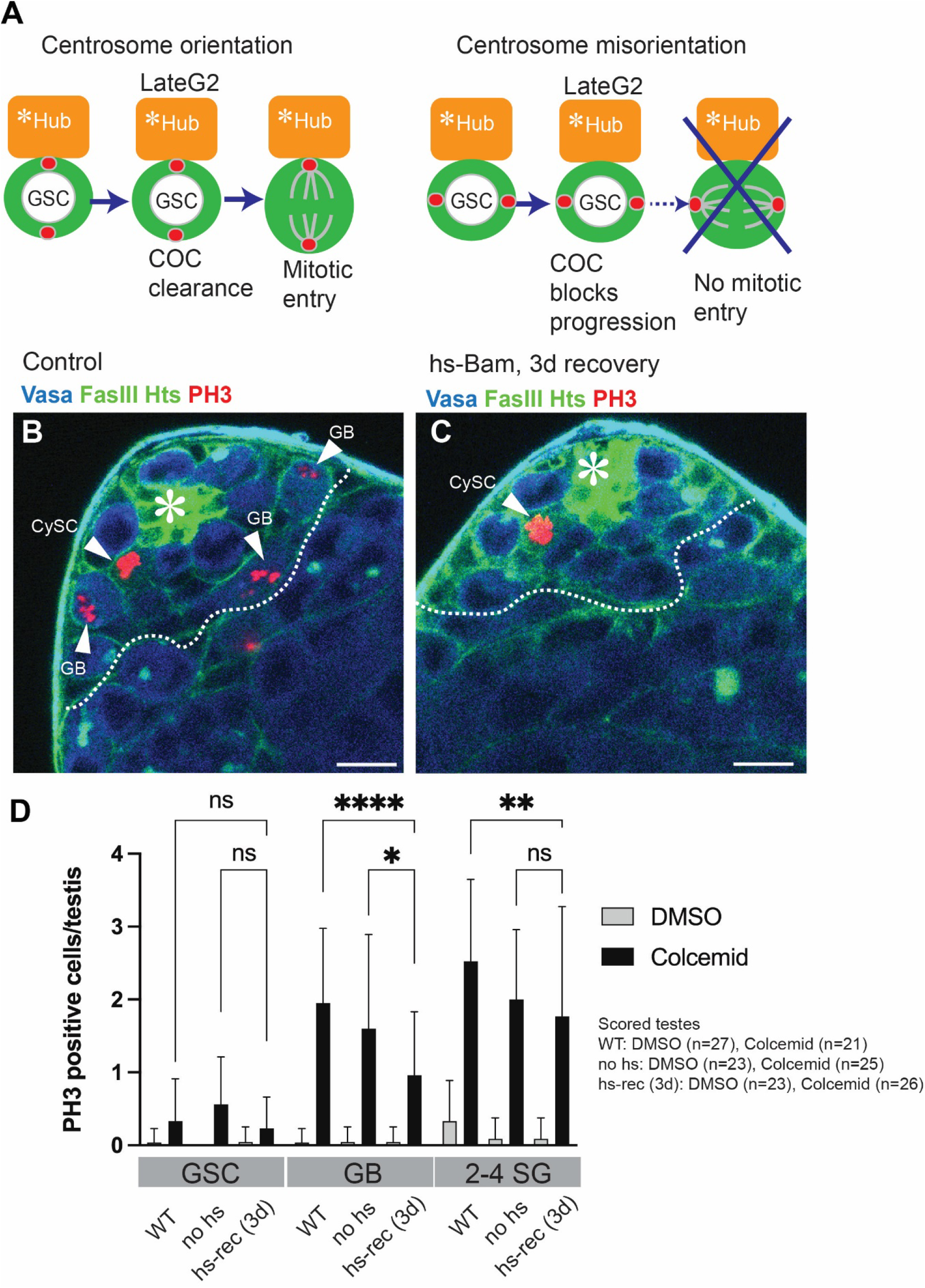
Centrosome orientation checkpoint is activated in dedifferentiating germ cells. A) Schematic of centrosome orientation checkpoint (COC). When centrosomes are positioned perpendicularly towards hub-GSC interface, COC is cleared in lateG2 phase and GSCs are allowed to enter mitosis. When centrosomes are mispositioned, COC blocks cells from progression to mitotic entry. *Hub=niche. **B, C**) Representative images of testes immunostained after 4.5-hour Colcemid treatment. White broken lines show approximate position of border between the GB population and SGs. Wild type testis (**B**) and hs-Bam testis (**C**) were used after 6-time heat-shock treatment and subsequent recovery for 3 days (see method for details). PH3 positive cells are indicated by white arrowheads. All scale bars represent 10 μm. The hub is marked by asterisks. **D**) PH3 positive cell number/testis after 4.5-hour Colcemid treatment. Testes were used at day-3 recovery point from forced differentiation of GSCs. P-values were calculated by Šídák’ s multiple comparisons test and provided as **** P < 0.00001, **P < 0.001, *P < 0.05 ns; non-significant (P≥0.05).

The COC has been shown to be active only in GSCs and not in differentiating GBs and SGs [30]. Since our observation indicated that dedifferentiating cells are likely in late G2 phase when they reattach to the hub, we hypothesized that the dedifferentiating cells may acquire COC and arrest in late G2 prior to niche reentry.

*Ex vivo* treatment of testes with microtubule depolymerizing drug Colcemid allow us to evaluate whether the cells has an active COC mechanism or not [30]. During the Colcemid treatment, cells without COC arrest at mitotic metaphase due to the spindle assembly checkpoint.

In contrast, cells with an active COC will arrest at late G2 phase. Therefore, we can assess the COC through counting phosphor-histone H3 (PH3) positive cells after 4.5-hour Colcemid treatment [30].

We depleted GSCs by hs-Bam expression then performed 4.5-hour Colcemid treatment and examine dedifferentiating cell population by looking at cells surrounding the niche (GBs) (Figure 3B, C). In early recovery phase, the cells closely located to the niche might be mixture of dedifferentiating cells and original GSCs just left from the niche. To avoid detecting original GSCs left from the niche, we used day-3 recovery phase when GSC number is almost recovered (7.85 ± 0.25, n=21). At day-3, the number of PH3 positive GBs was significantly lower (stopping at G2 phase) compared to the control as expected (Figure 3B-D).

It should be noted that we observed similar time between reattachment and 1^st^ mitotic entry in dedifferentiated cells even without GSC depletion (Figure S1, Table 2), further supporting the idea that COC arrested cells in the area surrounding the niche are likely dedifferentiating cells.

Taken together, our data strongly suggest that the dedifferentiating cells are arrested in COC, and this mechanism may act to ensure the correct division orientation of dedifferentiated cells after niche reentry.

### The checkpoint component, Baz is required for dedifferentiation

Activation of COC in dedifferentiating germ cells prior to niche attachment prompted us to hypothesize that the COC mechanism might be functionally linked to dedifferentiation, so that the system can ensure asymmetric division even in dedifferentiated cells. The polarity protein Bazooka/Par3 (Baz) is an essential component of the COC. Baz forms foci at the hub-GSC interface and serves as a docking site for the apical centrosome (Figure 4A). Baz-centrosome docking serves as an indicator of the correct centrosome orientation, and its phosphorylation likely triggers the release of the COC, allowing for GSC mitosis to occur [18].

**Figure 4.**
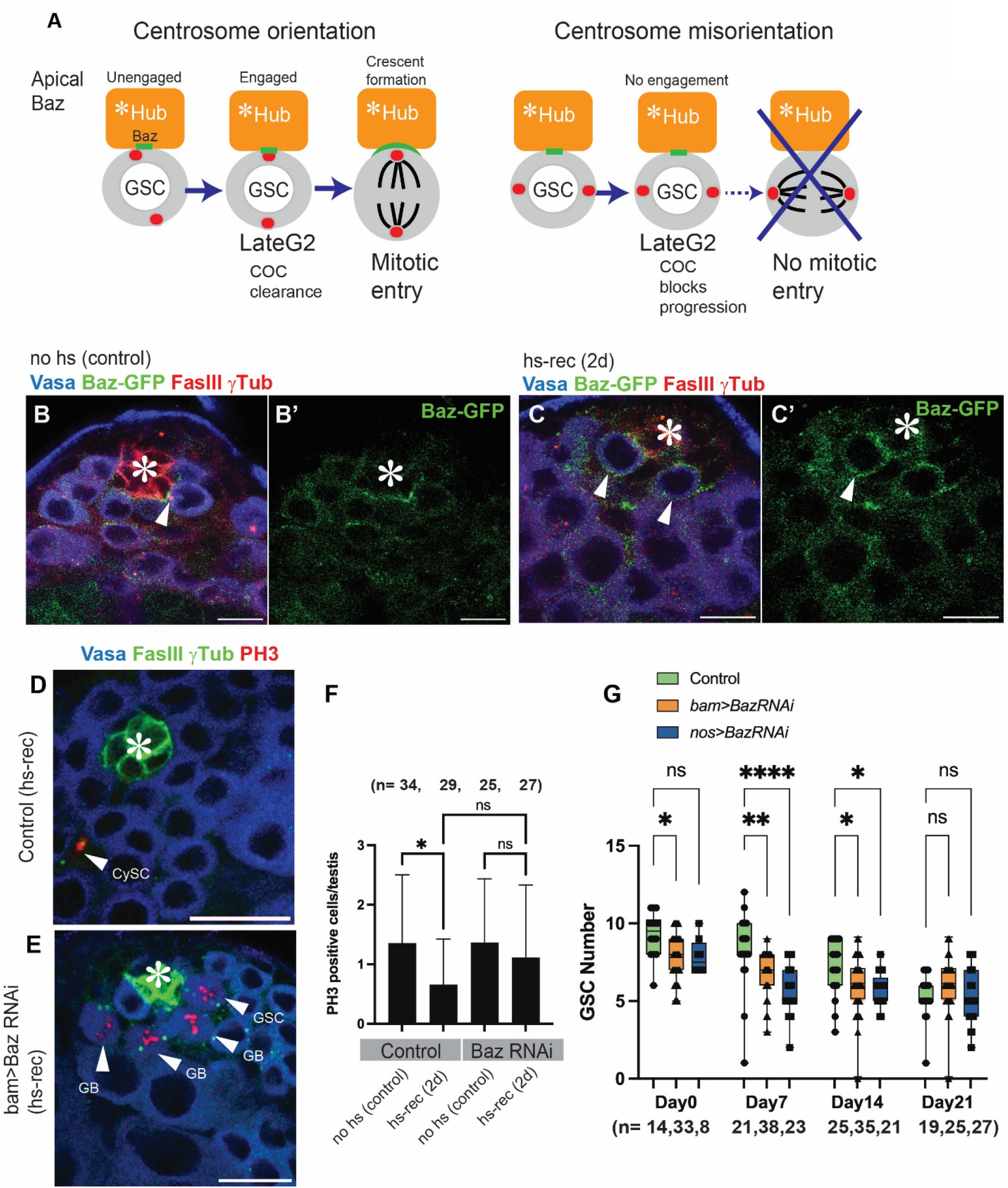
The checkpoint component, Baz is required for dedifferentiation. **A**) Schematic of Baz and centrosome interaction in COC. Apical centrosomes tightly interact (“engage”) with Baz patch at lateG2 phase, which triggers COC clearance. When centrosomes are mispositioned, centrosome-Baz docking/engagement does not occur, then COC sustains cells in lateG2 phase. *Hub=niche. **B, C**) Representative images of Baz-GFP trap testes with or without hs-Bam expression. Testes were immunostained at day-2 recovery point. PH3 positive cells are indicated by white arrowheads. **D, E**) Representative images of immunostained testes after 4.5-hour Colcemid treatment. Testes with or without Baz RNAi expression under the bamGal4 driver were used at day-2 recovery point from forced differentiation of GSCs. **F**) PH3 positive cell number/testis after 4.5-hour Colcemid treatment. Testes with or without Baz RNAi expression under the bamGal4 driver were used. P-values were calculated by Dunnett’ s multiple comparisons test and provided as * P < 0.05 or ns; non-significant (P≥0.05). **G**) Changes in GSC number during fly age in the flies of indicated genotypes. P-values were calculated by Šídák’ s multiple comparisons test and provided as **** P < 0.00001, ** P < 0.001, * P < 0.05, or ns; non-significant (P≥0.05).

To assess the requirement of COC activation for dedifferentiation to occur, we decided to determine the function of Baz in dedifferentiation. We first examined expression pattern of Baz protein in dedifferentiating germ cells. Baz-GFP trap signal is normally detected in any germ cells but especially concentrated in GSCs at the GSC-Hub interface. Particularly in G2-phase GSCs, we can observe Baz forms “patch-like” structure and capturing apical centrosomes (Figure 4A, B). In contrast, dedifferentiating/dedifferentiated GSCs often displays broad cortical pattern of Baz during early recovery phase (day2 after heat shock, Figure 4C), suggesting the possibility of Baz’ s involvement in mechanism of dedifferentiation.

We next knocked down Baz or expressed non-phosphorylatable Baz (Baz S151A, 1085A) in differentiating cells under the bamGal4 driver. After GSC depletion, we did not observe increased COC arrested GBs in Baz RNAi testes (Figure 4D-F), indicating that the Baz is required for G2 block in dedifferentiating cells, in addition to its known role on COC in GSCs [32]. Interestingly, we often observed PH3 germ cells are accumulating in the region surrounding the hub, indicating the possibility that dedifferentiating cells might be stuck near the niche due to the failure of their niche re-attachment (Figure 4E). Moreover, after forced depletion of GSCs, the recovery of GSCs was significantly slower and incomplete both in Baz RNAi and Baz SA-expressing testes compared to the control testes (Figure S4). In addition to the defect of dedifferentiation, we observed a significant reduction in the number of GSCs in both the nosGal4 and bamGal4-driven Baz-RNAi flies (Figure 4G), indicating that Baz is required for GSC number maintenance through dedifferentiation. These data indicate that the Baz-mediated G2 arrest is required for dedifferentiation to occur.

Taken together, this study provides clear evidence that a dedifferentiated stem cell can divide asymmetrically. Moreover, the functional link between dedifferentiation and COC allows only cells with activated COC to reenter the niche to ensure correctly oriented spindles upon niche reentrance.

## Discussion

In this study, we show that the polarity checkpoint, centrosome orientation checkpoint (COC), is activated in dedifferentiating germ cells prior to niche reentry. The checkpoint component, Baz is required for maintenance of GSC number through dedifferentiation, indicating that dedifferentiation is mechanistically linked to COC, thus ensuring asymmetric division even in dedifferentiated stem cells.

Dedifferentiated cells can be marked by lineage tracing method to permanently mark cells once experienced Bam positive linage [8, 9]. However, this method is not sensitive enough to detect majority of dedifferentiated cells as previous work showed that the GB to four-cell SGs are most potent for dedifferentiation using live imaging [7]. Therefore, we used similar method to monitor behavior of dedifferentiated cells directly.

During asymmetric cell division, certain cellular components are distributed unevenly between the daughter cells. For example, the mother centrosome is always positioned at the hub-GSC interface and is passed on to the GSC [31]. Additionally, old, and new histone H3 and sister chromatids of the X and Y chromosomes are distributed asymmetrically during the division of male GSCs [33, 34]. These observations indicate that there are inherent differences between the two daughter stem cells. When asymmetric outcome is compromised through dedifferentiation, asymmetric inheritance of intrinsic factors can be compromised, raising the question of whether dedifferentiated GSCs behave in the same way as original GSCs.

Nonetheless, our study demonstrates the remarkable capacity of dedifferentiated cells to reacquire asymmetric division in the niche. We propose that the functional link between dedifferentiation and stem-cell specific checkpoint ensures asymmetric division of stem cells after reentrance to the niche. In summary, the present study provides a new insight into how asymmetric division is ensured in dedifferentiated stem cells.

## Materials and methods

### Fly husbandry and strains

Flies were raised on standard Bloomington medium at 25°C (unless temperature control was required). 0-to 7-day-old adults were used for all experiments unless specific ages of flies are noted. The following fly stocks were obtained from Bloomington stock center (BDSC); nosGal4 (BDSC64277)[35]; Fly-FUCCI, pUASp-GFP.E2f1.1-230, UASp-mRFP1.CycB.1-266 (BDSC 55100), hs-bam X (BDSC24636) or 3^rd^ (BDSC24637); yw (BDSC189) was used for wildtype. UAS-Baz-S151A S1085A [36], bamGal4 on 3rd Chromosome was kind gift from Yukiko Yamashita. Baz-GFP (Fry-trap, CC01941), [37, 38], bamGal4VP16 on X Chromosome was kind gift from Michael Buszczak.

### Induction of dedifferentiation

A previously established technique was used to experimentally induce dedifferentiation with slight modifications [6]. Briefly, 0-3 days old flies raised at room temperature with one allele of the hsBam transgene on either the X(BDSC24636) or 3^rd^(BDSC24637) chromosome were subjected heat-shock treatment by immersing the fly vials with fly food in a 37 degree water bath for 30 mins, twice a day, for a total of six times. The flies were incubated at 29 degrees in between consecutive heat-shock treatments and moved to room temperature after the last heat-shock for recovery. The testes were then dissected after the desired time of recovery.

### Live Imaging

Live imaging was performed following previously established technique [39]. The testes were dissected into 1X Becker Ringer’ s solution [39] and then mounted onto a 35mm Glass Bottom Dishes (Nunc). 200 μL of 1 mg/mL poly-L-lysine (Sigma) was pipetted onto the coverslip portion of the imaging dish and incubated for 5-to 7-hours at room temperature. Then, poly-L-lysine solution was replaced to the Becker Ringer’ s solution and testes were mounted onto poly-L-lysine layer with the tip of the testes oriented toward bottom. Next, Becker Ringer’ s solution was slowly removed and pre-warmed (room temperature) Schneider’ s Drosophila medium containing 10% fetal bovine serum and glutamine–penicillin–streptomycin (Sigma) was added to the dish.

Z-stacks (1mm interval, for 10∼15 stacks) were taken using a Zeiss LSM800 airyscan with a 63× oil immersion objective (NA = 1.4) every 5 to 10 minutes for 16 hours. Preset tiling function was used for automated acquisition.

### Colcemid treatment

Colcemid treatment was performed as described previously [30]. Testes were dissected and transferred to Schneider’ s Drosophila medium (Gibco) containing 10% fetal calf serum (Gibco) and glutamine–penicillin–streptomycin (Sigma). Colcemid (Calbiochem) was added at a final concentration of 100 μM. Same volume of DMSO was added for control samples. Samples were incubated on nutator at room temperature for 4.5 h, then processed for Immunofluorescence staining.

### Immunofluorescence staining

Testes were dissected in 1x phosphate-buffered saline (PBS) (Fisher Scientific) and fixed in 4% formaldehyde (Electron Microscopy Sciences) in PBS for 30–60 minutes. Next, testes were washed in PBS with 0.2% TritonX-100 (Fisher Scientific) (PBST) for at least 60 minutes, followed by incubation with primary antibody in 3% bovine serum albumin (BSA) (Sigma) in PBST at 4°C overnight. Next day, samples were washed for 60 minutes (three times for 20 minutes each) in PBST, incubated with secondary antibody in 3% BSA in PBST at room temperature for 2 hours and then washed for 60 minutes (three times for 20 minutes each) in PBST. Samples were then mounted using VECTASHIELD with 4’, 6-diamidino-2-phenylindole (DAPI) (Vector Lab).

The primary antibodies used were as follows: rat anti-Vasa (1:20; developed by A. Spradling and D. Williams, obtained from Developmental Studies Hybridoma Bank (DSHB); mouse anti-hu-li tai shao (Hts) (1:20, 1B1; DSHB); mouse-anti-Fasciclin III (FasIII) (1:20, 7G10; DSHB); mouse anti-γ-Tubulin (1:400, GTU-88; Sigma-Aldrich); rabbit anti-phosphorylated (Thr3) histone H3 (1:200; clone JY325, Upstate). AlexaFluor-conjugated secondary antibodies (Abcam) were used at a dilution of 1:400.

### Statistical analysis and graphing

No statistical methods were used to predetermine sample size. The experiment values were not randomized. The investigators were not blinded to allocation during experiments and outcome assessment. All experiments were independently repeated at least 3 times to confirm the results. Statistical analysis and graphing were performed using GraphPad prism9. All data are shown as means ± s.d. The adjusted p values are provided; shown as *P<0.05, **P<0.01, ***P<0.001, ****P<0.0001; NS, non-significant (P≥0.05).

## Acknowledgements

We thank Yukiko M. Yamashita, Michael Buszczak, Andreas Wodarz and the Bloomington *Drosophila* Stock Center and the Developmental Studies Hybridoma Bank for reagents, Erika Matunis for valuable suggestion for live imaging. This research is supported by R35GM128678 from National Institute for General Medical Sciences and start-up funds from UConn Health (to M.I.).

## Author Contributions

M.I., M.B.B. and N.P. conceived the project. M.I., M.B.B., A.T., R.C. designed and executed experiments and analyzed data. M.B.B. and M.I. drafted manuscript. All authors edited the manuscript.

