## Supplemental Figures for "Dedifferentiating germ cells regain stem-cell specific polarity checkpoint prior to niche reentry"

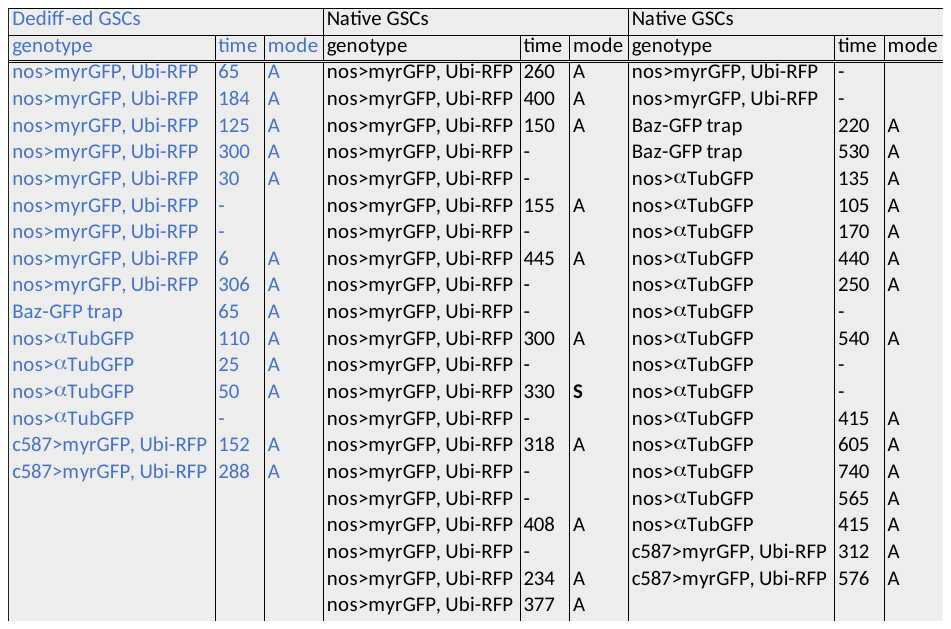

**Table 1. Division of dedifferentiated GSCs native GSCs after forced depletion of GSCs.**

Indicated genotypes combined with hs-bam were used. Time lapse imaging was performed at 32 to 48 hour- recovery period after hs-Bam expression. Native GSCs are defined as the GSCs that were already attached to the hub when imaging was started. For dedifferentiated cells, “time” means the minutes from the hub reattachment to the entry of 1^st^ mitosis (nuclear envelope break down; NEBD). For native GSCs, “time” indicates the minutes taken to the entry of 1^st^ mitosis after the imaging was started. When no division was observed, the time column shows (-). “mode”; division mode of the cell was categorized to be A or S (A: asymmetric division, S: symmetric division) based on the orientation of the cell division.

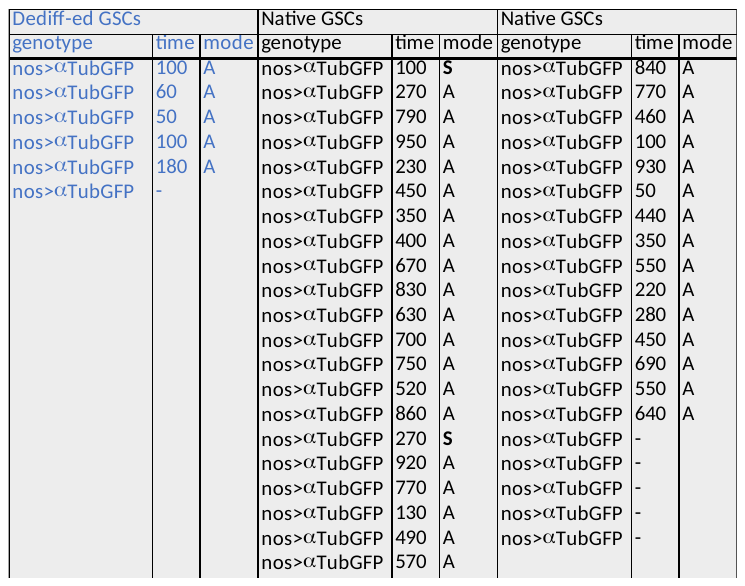

**Table 2. Tracing of dedifferentiated GSCs and native GSCs without depletion of GSCs.**

Flies expressing αTub-GFP under the control of nosGal4 driver were used. Native GSCs are defined as the GSCs that were already attached to the hub when imaging was started. For dedifferentiated cells, “time” means the minutes from the hub reattachment to the entry of 1^st^ mitosis (nuclear envelope break down; NEBD). For native GSCs, “time” indicates the minutes taken to the entry of 1^st^ mitosis after the imaging was started. When no division was observed, the time column shows (-). “mode”; division mode of the cell was categorized to be A or S (A: asymmetric division, S: symmetric division) based on the orientation of the cell division.

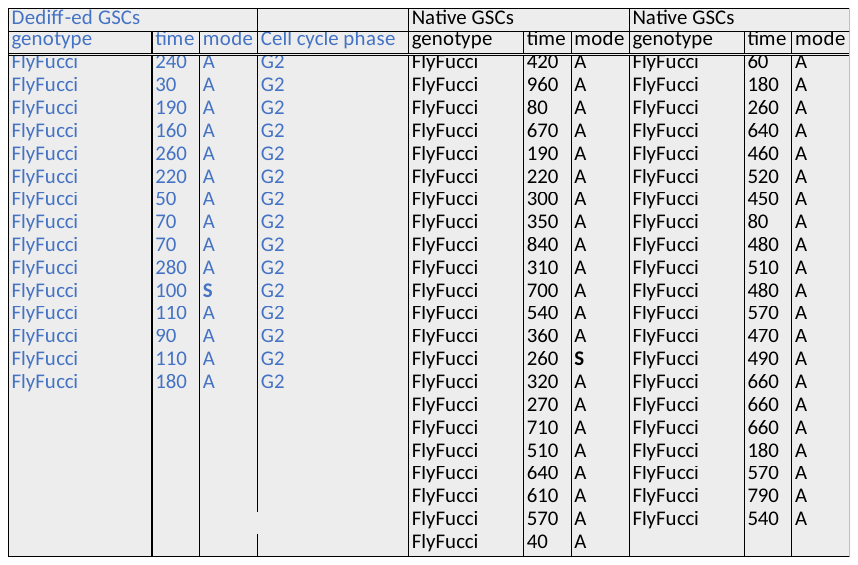
**Table 1. Tracing of dedifferentiated GSCs and native GSCs expressing fly-FUCCI cell cycle marker.**  Flies expressing fly-FUCCI marker under the control of nosGal4 driver combined with hs-bam were used. Time lapse imaging was performed at 32 to 48 hour- recovery period after hs-Bam expression. Native GSCs are defined as the GSCs that were already attached to the hub when imaging was started. For dedifferentiated cells, “time” means the minutes from the hub reattachment to the entry of 1^st^ mitosis (nuclear envelope break down; NEBD). For native GSCs, “time” indicates the minutes taken to the entry of 1^st^ mitosis after the imaging was started. When no division was observed, the time column shows (-). “mode”; division mode of the cell was categorized to be A or S (A: asymmetric division, S: symmetric division) based on the orientation of the cell division. Cell cycle phase was judged by fly-FUCCI (see main text).

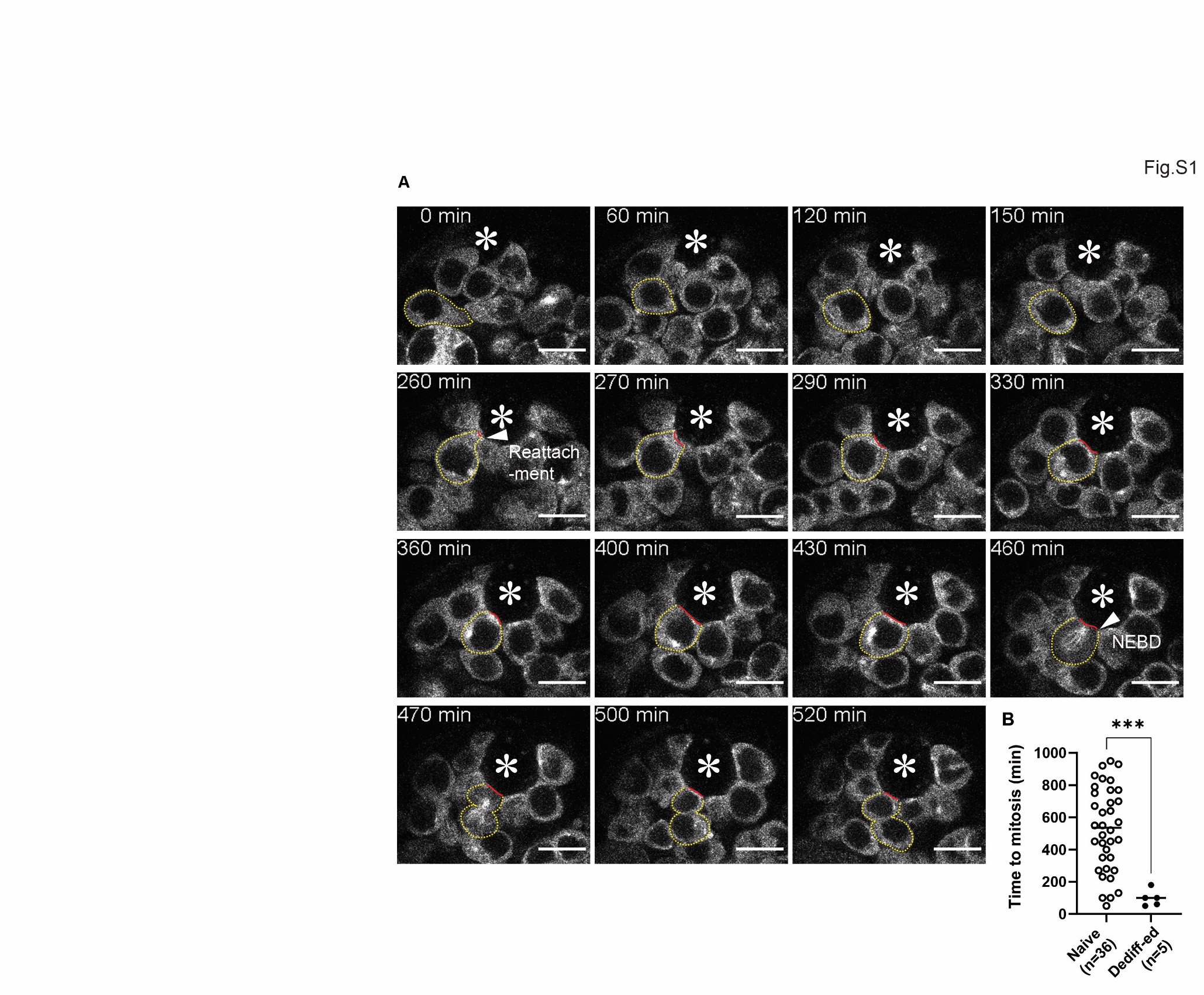

**Figure S1. Dedifferentiated GSCs divide immediately upon niche reattachment.**

**A**) Representative time-lapse live images of a dedifferentiation event without depletion of GSCs. Germ cells are visualized by expression of UAS-αTubGFP under the control of nosGal4 driver. A dedifferentiating germ cell is encircled by yellow dotted lines. The interface of dedifferentiated GSC and hub cell is marked by red lines. NEBD=nuclear envelope breakdown. **B**) The time taken between hub- reattachment and 1^st^ mitotic division (NEBD) for all recorded dedifferention events listed in Table 3 is plotted to the graph. The cases of “not divided” were not included in the graph. P-values were calculated by a 2-tailed student-t-test and provided as *** P < 0.0001. “n” indicates the number of scored GSCs. Horizontal lines in the graph represent mean values. All scale bars represent 10 μm. The hub is marked by asterisks.

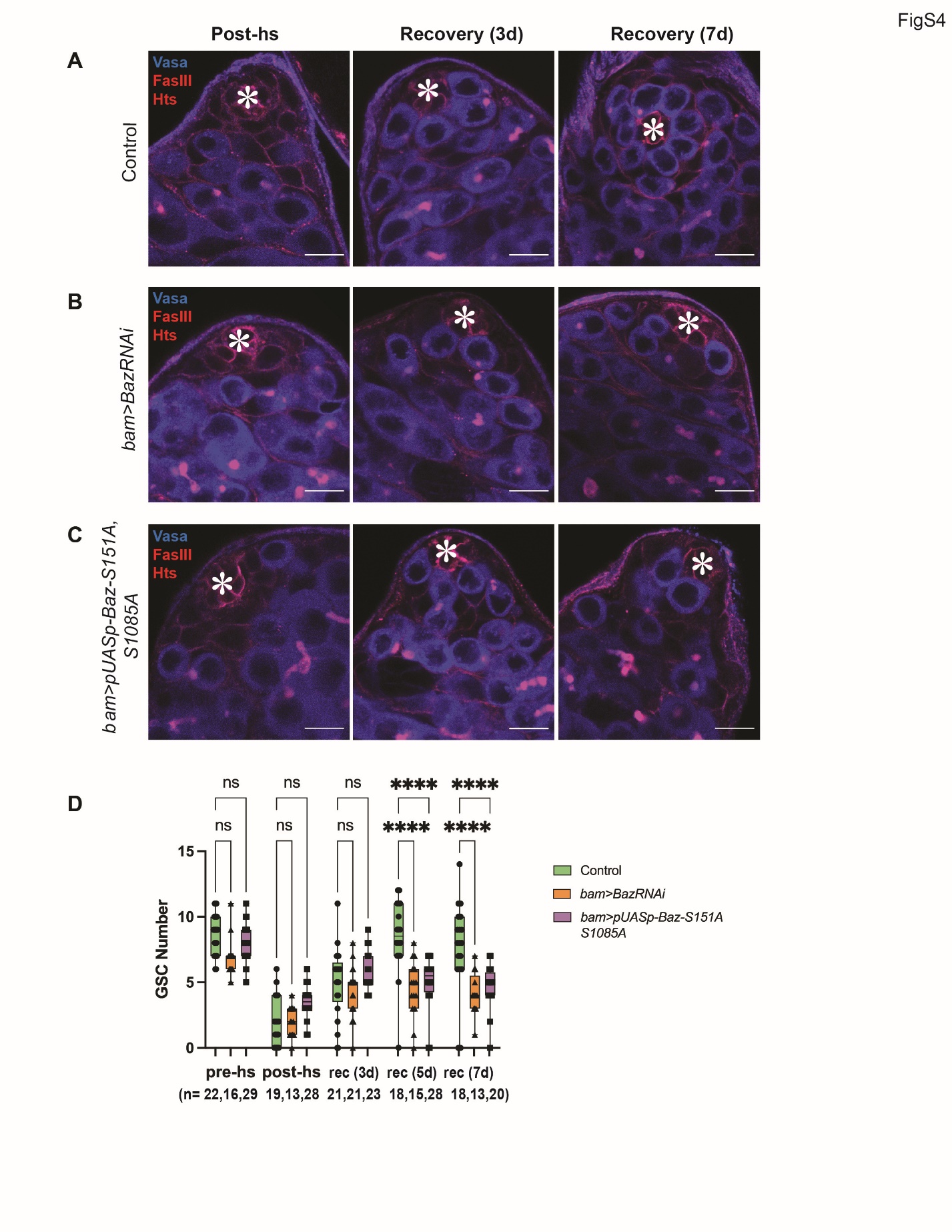

**Figure S4.** **Baz is required for dedifferentiation.**

**A-C**) Representative images of testis tip after depletion of GSC by expressing Bam (post HS; after 6-time heat shock treatment) and after 3-day recovery in room temperature culture without (**A**) or with expression of Baz RNAi (**B**) or phosphorylation defective Baz (UAS-BazS151A, S1085A, **D**). Changes in GSC number during recovery from forced differentiation of GSCs without or with expression of Baz RNAi or phosphorylation defective Baz (UAS-BazS151A, S1085A. P-values were calculated by Šídák's multiple comparisons test and provided as **** P < 0.00001 or ns; non-significant (P≥0.05). All scale bars represent 10 μm. The hub is marked by asterisks.
